# CoDEX2: full-spectrum copy number variation detection by high-throughput DNA sequencing

**DOI:** 10.1101/211698

**Authors:** Yuchao Jiang, Rujin Wang, Eugene Urrutia, Ioannis N. Anastopoulos, Katherine L. Nathanson, Nancy R. Zhang

**Affiliations:** Department of Biostatistics, Gillings School of Global Public Health, University of North Carolina, Chapel Hill, NC 27599, USA.; Department of Genetics, School of Medicine, University of North Carolina, Chapel Hill, NC 27599, USA.; Lineberger Comprehensive Cancer Center, University of North Carolina, Chapel Hill, NC 27599, USA; Division of Translational Medicine and Human Genetics, Department of Medicine, Perelman School of Medicine, University of Pennsylvania, Philadelphia, PA 19104, USA.; Abramson Cancer Center, Perelman School of Medicine, University of Pennsylvania, Philadelphia, PA 19104, USA.; Department of Statistics, The Wharton School, University of Pennsylvania, Philadelphia, PA 19104, USA.

**Keywords:** copy number variation, normalization, next-generation sequencing, latent factor, negative control

## Abstract

High-throughput DNA sequencing enables detection of copy number variations (CNVs) on the genome-wide scale with finer resolution compared to array-based methods, but suffers from biases and artifacts that lead to false discoveries and low sensitivity. We describe CODEX2, a statistical framework for full-spectrum CNV profiling that is sensitive for variants with both common and rare population frequencies and that is applicable to study designs with and without negative control samples. We demonstrate and evaluate CODEX2 on whole-exome and targeted sequencing data, where biases are the most prominent. CODEX2 outperforms existing methods and, in particular, significantly improves sensitivity for common CNVs.

## Background

Copy number variations (CNVs) are large deletions and duplications of segments of the chromosome. CNVs are pervasive in the human genome and play a causal role in diseases such as cancer [1]. In the study of disease, CNVs usually appear in two contexts: *germline* CNVs refer to inherited variants, many of which are polymorphic at the population level [2]; in contrast, *somatic* CNVs, also referred to as copy number aberrations (CNAs), are the copy number changes resulting from somatic mutations, such as those commonly observed in cancer. Germline CNVs can also be described as *common* or *rare* based on their population frequencies. This paper addresses the problem of detection of both germline and somatic CNVs, and, in particular, of improving detection sensitivity for common CNVs in both categories.

With the dramatic growth of sequencing capacity and the accompanying drop in sequencing cost, massively parallel next-generation sequencing offers appealing platforms for genome-wide CNV detection. Whole-genome sequencing (WGS) offers an unbiased genome-wide approach to detect CNVs, while whole-exome sequencing (WES) and targeted sequencing allow the identification of disease-associated variants in coding regions with direct functional interpretation. Despite the rapid technological progress, CNV detection using high-throughput sequencing still faces analytical challenges due to the pervasive biases and artifacts that are introduced during library preparation and sequencing. Proper data normalization is crucial, especially for WES and targeted sequencing, where technical biases are usually magnitudes larger than CNV signal.

For studies in which deep coverage of specific genome regions is desired [3–5], or where the cohort of interest is large, WES and targeted sequencing are often preferred as cheaper alternatives to WGS. For example, the DiscovEHR Collaboration (http://www.discovehrshare.com) has sequenced the exomes of more than 50,000 participants, and the Exome Aggregation Consortium (ExAC; http://exac.broadinstitute.org) has generated aggregated WES data for 60,706 unrelated individuals taken from multiple disease-specific and population genetic studies. This paper was primarily motivated by the challenges in WES and targeted sequencing data, but the methods developed here have also been integrated into a CNV detection pipeline for WGS data [6], where it is shown that the normalization methods prescribed here allow for more accurate CNV calls using Mendelian concordance as metric.

CNV detection (by WES and targeted sequencing) is primarily based on the detection of local changes in read coverage along the genome. This analysis scheme is based on the simple intuition that regions with copy number gain should have increased read coverage, and regions with copy number loss should have decreased read coverage. However, read coverage depends not just on copy number but also on many other factors, such as GC content [7], mappability [8], and other local sequence characteristics [9]. Therefore, read coverage tends to fluctuate even when there is no CNV, and is especially variable in WES and targeted sequencing data due to the biases and artifacts introduced during the targeting and amplification steps [10–12].

Many methods are available for CNV detection in high throughput sequencing data [10–16]. Despite much progress however [17], significant challenges still remain. Independent benchmark results from multiple studies [10, 12, 14, 18] show that existing methods suffer from low precision and recall rates, especially for the detection of common germline CNV signals. These results are not unexpected, since in WES and targeted sequencing data, and also to an extent in WGS data, the technical background bias for each genomic target varies across samples, leading to false positives and negatives if not properly removed. To remove this technical background, recent methods have relied on low-dimensional linear factor models to capture the background bias [10, 11], which tend to control for false positives. However, these low-dimensional linear factor models tend to remove common CNVs that correlate with the estimated factors. CLAMMS [19] is developed to recover common CNV signals by WES but is not suitable for cancer samples, where recurrent somatic copy number changes are prominent. In this paper, we demonstrate that this issue also affects the detection of somatic CNVs in cancer genomes, as CNVs that are recurrent across multiple cancer samples can be accidentally removed in the normalization step. Due to these limitations, current genetic studies using WES and targeted sequencing data have focused mostly on single nucleotide variations and, at best, rare CNVs [20–22].

Herein, we propose CODEX2 for full-spectrum CNV detection in high-throughput sequencing data. In this context, full-spectrum implies the sensitive detection of both common and rare CNVs. CODEX2 can be applied to two scenarios: when there are ‘cases’ and ‘controls’ and the goal is to detect CNVs that are enriched in the case samples (Figure 1A); and when control samples are lacking and the goal is simply to profile all CNVs in all samples (Figure 1B). We evaluated CODEX2 in three ways. First, we reanalyzed the WES data of the HapMap samples from the 1000 Genomes Project [2], with matched microarrays and experimental validation [23–25] to assess performance. Our results demonstrate that CODEX2 significantly improves both sensitivity and specificity over existing methods, especially for common CNVs. Next, we applied CODEX2 to targeted sequencing data of melanoma cancer cell lines, patient derived xenografts (PDX) and tumor biopsies, and successfully identified recurrent CNVs whose frequencies are highly concordant with those obtained from a separate cohort studied by the Cancer Genome Atlas (TCGA) [1]. Finally, we performed extensive simulations to benchmark existing methods and to explore how key variables, such as population frequency and CNV length, influence performance. CODEX2 is compiled as an open-source R-package available at https://github.com/vuchaoiiang/CODEX2.

**Figure 1.**
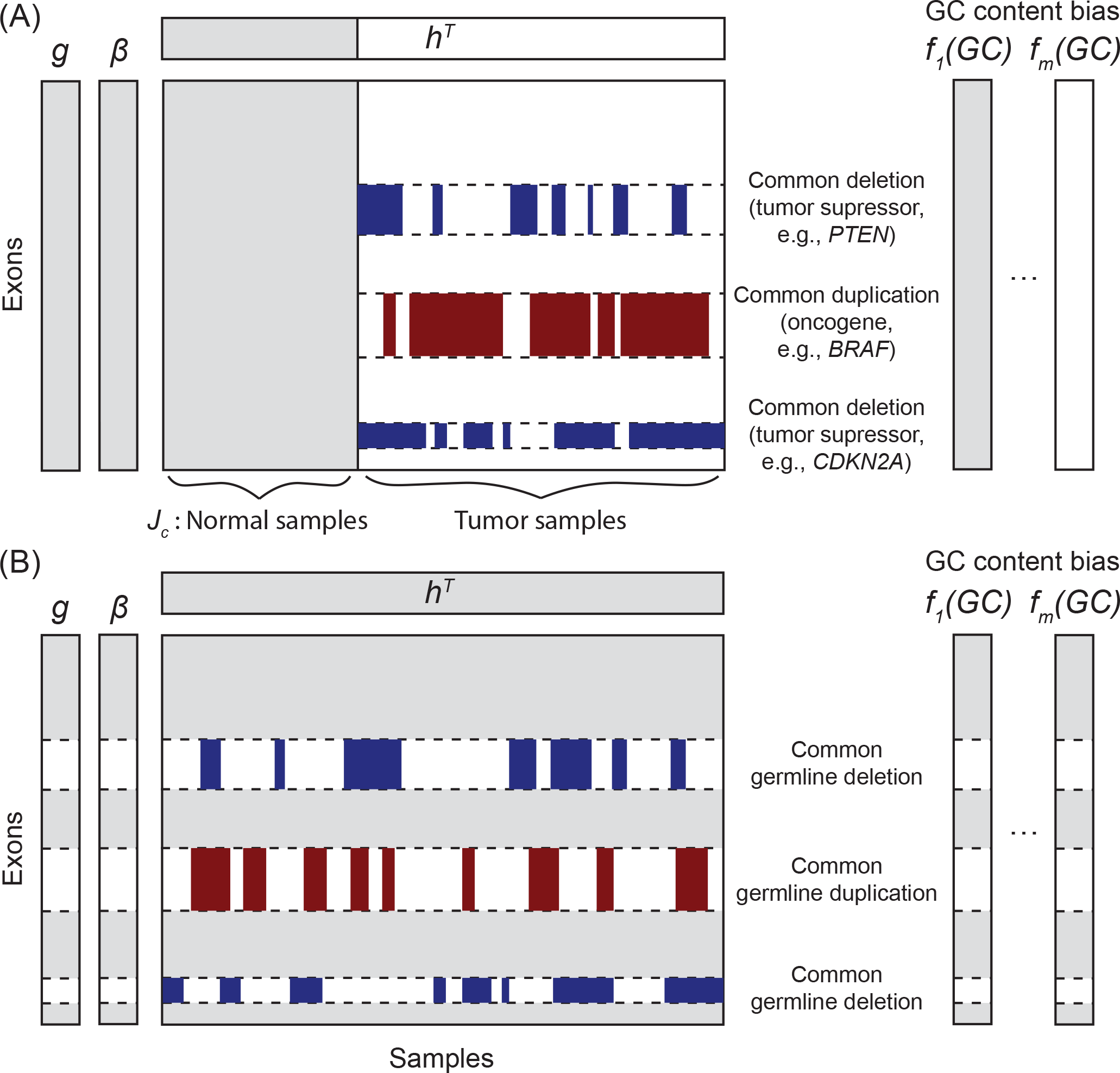
CODEX2 can be applied under two experimental designs to identify common and rare, germline and somatic CNVs. (A) CODEX2 with negative control samples. The goal is to recover both rare and common CNV signals present disproportionately in the cases versus the controls. (B) CODEX2 without negative control samples. The goal is to identify all CNVs in all samples, e.g., to detect germline CNVs in healthy individuals without prior knowledge of disease status.

## Results

### Methods overview

Figure 1 illustrates the two experimental designs for which CODEX2 can be applied: (i) case-control design with a group of negative control samples, where the goal is to detect CNVs disproportionately present in the ‘cases’ versus the ‘controls’ (Figure 1A); and (ii) detection of all CNVs present in all samples design, such as in ExAC (Figure 1B). The key innovation in CODEX2 is the way that it harnesses *negative control genome regions and/or negative control samples* in its genome-wide latent factor model for sample- and position-specific background correction. The negative control genome regions defined by CODEX2 are regions that do not harbor *common* CNVs, but that are still allowed to harbor rare CNVs, and can be constructed from existing studies or learned from data.

Figure 2 illustrates how CODEX2 normalization improves upon CODEX and SV-based normalization methods such as XHMM [11]. For simplicity, and without loss of generality, we represent the background bias by a 1-dimensional latent factor, which we call the "latent batch effect”. Against this background, we further assume that there are two CNVs: Region A, in which the carrier status (the vector indicating whether each sample is a carrier) is correlated with the underlying latent batch, and region B, in which the carrier status is uncorrelated with the latent batch. The signal for these two CNV regions is obscured by the background batch effect in the observed data. In a standard SVD or CODEX normalization, which does not make use of negative controls, all samples and all genomic regions are used in the estimation of the latent background model, resulting in contamination of background estimates by the CNV signal. The contamination is especially severe for CNV A, which is correlated with the one-dimensional background batch. For CODEX2, in scenarios for which negative controls are available, only the negative controls are used to fit the latent factor model, which is then used to predict the background bias of the rest of the samples. In scenarios for which negative control samples are not available but negative control regions are identified, the latent factor model is fitted only using the negative control regions, and then used to “fill in” the background bias for the rest of the regions. In this way, we avoid the contamination of the background estimates by the CNV signal, thus attaining better separation of signal from noise, as can be seen from the histograms of the normalized *z*-scores. More details are given in Methods.

**Figure 2.**
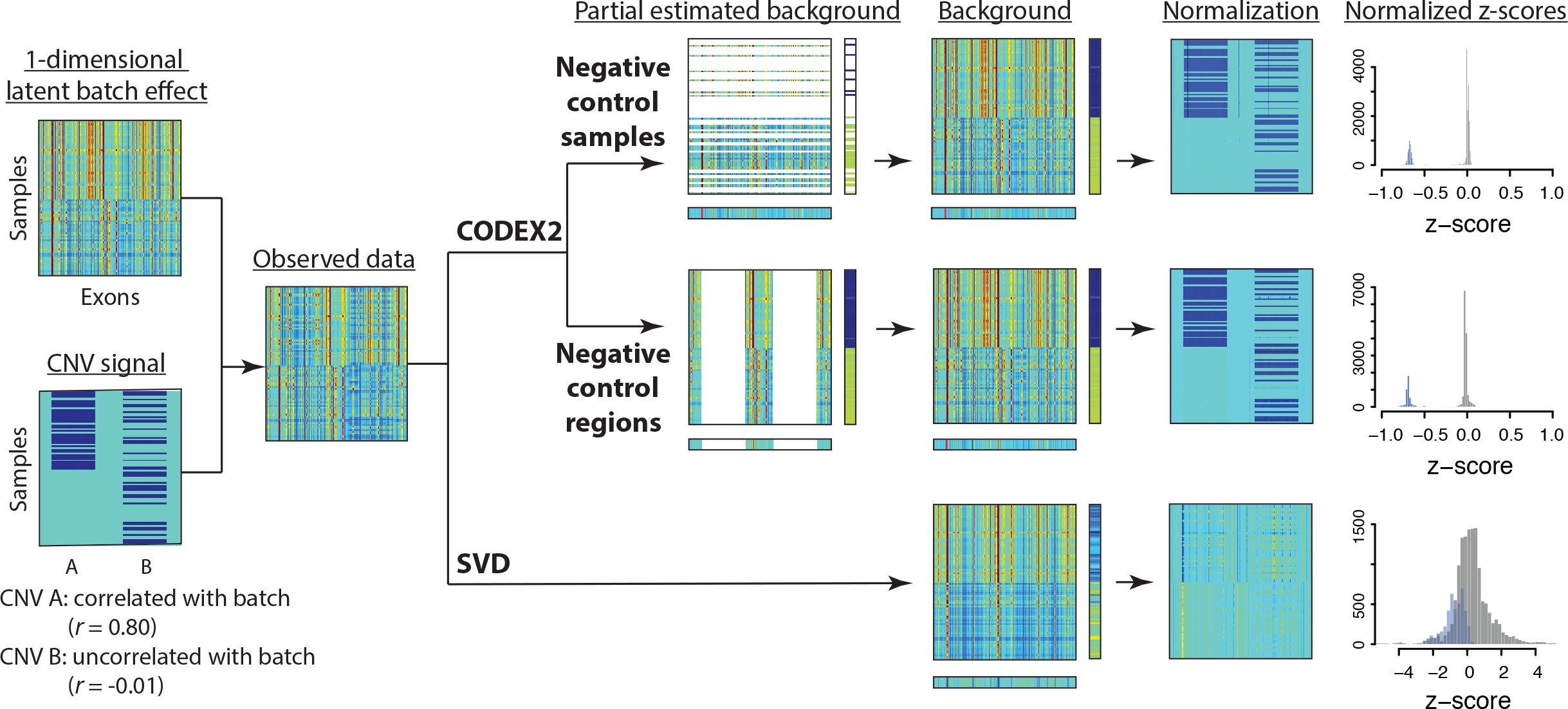
Outline of analysis framework with benchmark against SVD-based methods. Results are based on *in silico* simulations where the ground truth with one dimensional latent factor is known. One CNV signal is correlated with batch effect; the other is not. CODEX2 utilizes negative control samples and negative control regions and outperforms SVD-based methods.

CODEX2 estimates a separate background factor for each genomic target/region in each sample, which can then be used to normalize the observed coverage and detect CNV regions using the recursive Poisson-likelihood segmentation algorithm in Jiang et al. [12].

### Analysis of WES and WGS data from the 1000 Genomes Project

We first evaluated CODEX2 by reanalyzing a publicly available WES data set from the 1000 Genomes Project [2], which contains 90 healthy individuals (Supplementary Table 1). Forty individuals are Utah residents with northern and western European ancestry; 20 are Japanese from Tokyo; 26 are Yoruba people from Ibadan. Gender is well balanced with 44 males and 46 females. The dataset contains two batches that are sequenced at the Baylor College of Medicine and the Washington University Genome Sequencing Center, respectively.

To assess performance of CODEX2 and to benchmark against existing methods, we used WGS CNV calls from phase 3 release [2], as well as the experimentally validated CNVs from three previous microarray studies [23–25] as gold standards. Specifically, we adopted stringent quality control procedures (i.e., the reported CNV must overlap with at least two and at most 20 exons, have less than 5% NA rate across all samples, and have at most three copy number states). These ‘gold-standard’ CNVs, shown in Supplementary Table 2, were categorized as common if they are present in more than 10% of the samples, and rare otherwise. Using these ‘validation sets’, we were able to assess the performance of CLAMMS [19], XHMM [11], EXCAVATOR [14], CODEX [12], and CODEX2 to detect common and rare CNVs, with results shown in Supplementary Table 3. Figure 3 and Supplementary Table 4 show the precision and recall rates across the four benchmarked methods, using the CNVs validated by each of the four studies as ground truth. The grey lines are the contours of the *F*-measure, defined as the harmonic mean of precision and recall. XHMM, which is designed for detection of rare CNVs, lacks sensitivity to common CNVs, so does EXCAVATOR. CODEX detects proportionately more common CNVs but still lacks sensitivity. CLAMMS retains high precision rates overall but suffers from low sensitivity. CODEX2 achieves a recall rate of 92.8%, 60.7%, 79.2%, and 66.2% in the four validation sets, respectively, while simultaneously making substantial improvements to specificity. CODEX2 does not dominate in the phase 3 WGS validation set, potentially due to the false calls in the set (Supplementary Figure 1). Overall, CODEX2 achieves good performance for both common and rare CNVs, with significant improvement in precision and recall for common CNV detection.

**Figure 3.**
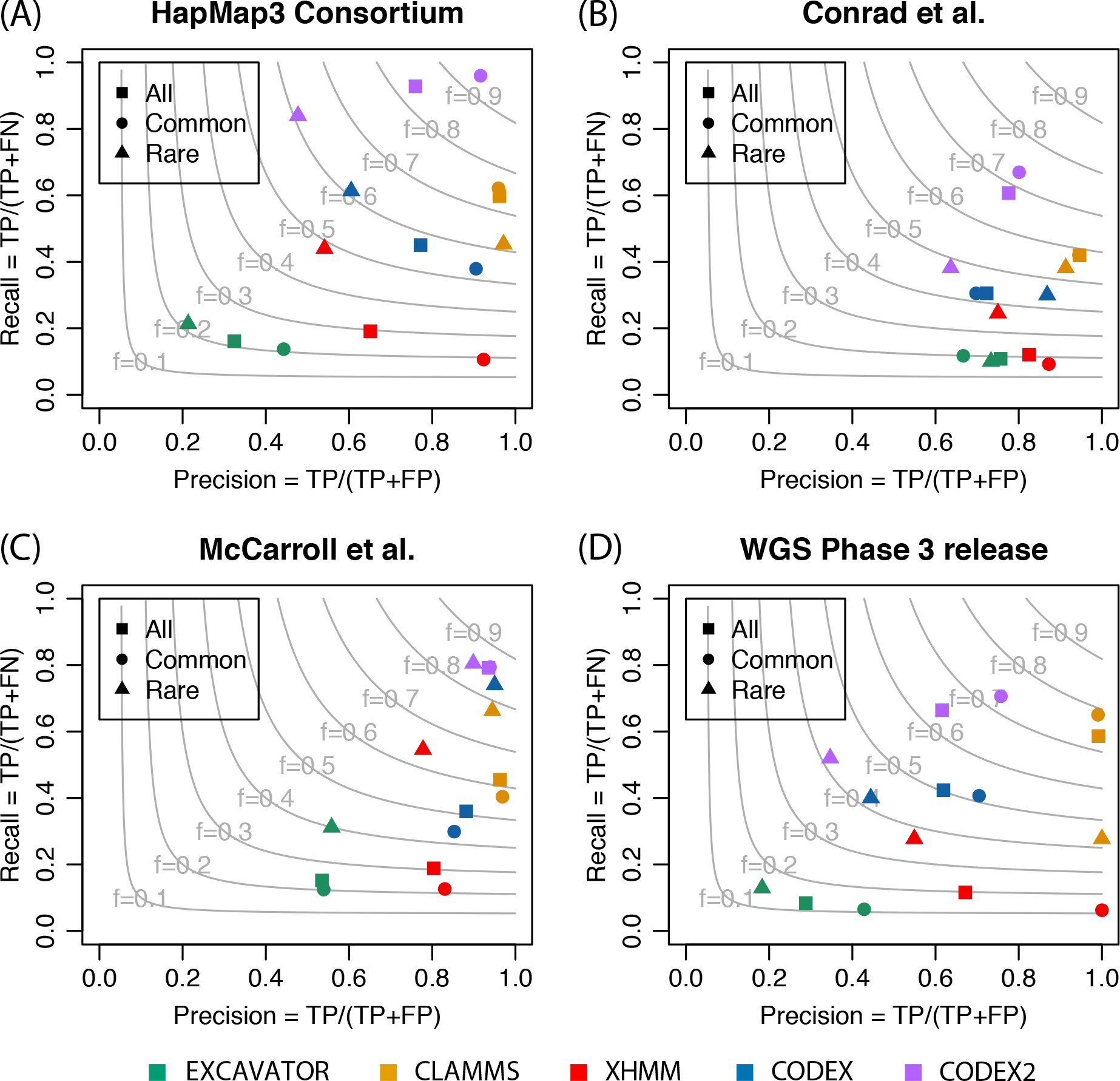
Assessment of CNV calls on WES data from the 1000 Genomes Project by microarray and WGS calls. CNV calls by XHMM, EXCAVATOR, CODEX and CODEX2 are validated against genotyping calls from (A) HapMap 3 Consortium [25], (B) Conrad et al. [24], and (C) McCarroll et al. [23], as well as WGS CNV calls from the phase 3 release [2]. Grey contours show *F*-measures as harmonic means of precision and recall rates. CODEX2 has significantly improved precision and recall for common CNVs and has the highest *F*-measure among all methods.

We further demonstrate CODEX2 on a WGS dataset of 108 individuals from the 1000 Genomes Project (Supplementary Table 1), out of which there are three family trios (Supplementary Table 5). We applied CODEX2 and CNVnator [26] to profile germline CNVs and assessed performance using the Mendelian concordance rates between children and their parents as metric. CODEX2 and CNVnator on average returned 1011 and 118 CNVs per individual, respectively (see Supplementary Table 6 for calling results). CODEX2 has higher number of overlapped CNVs between children and parents (Supplementary Figure 1) as well as higher call set quality (Supplementary Figure 2). Specifically, sequencing data, especially WES, are known to contain reproducible artifacts that lead to false positives shared between samples. Thus, the degree of overlap in CNV calls between samples is, per se, not a good measure of accuracy; In fact, if no normalization is performed, the CNV calls would comprise mostly of false positives and would have an inflated cross-sample concordance close to one. Therefore, we assess call accuracy using the ratio of child-parent concordance over the concordance between the two unrelated parents. This measure of concordance enrichment more accurately reflect call set quality.

### Analysis of targeted sequencing data of melanoma cases and controls

We further applied CODEX2 to a cohort study of melanoma from Garman et al. [27] including 334 cases (untreated human melanoma cell lines, patient-derived xenografts, and tumors) and 16 controls. Samples were sequenced on a custom capture panel of 108 genes previously implicated in tumorigenesis of melanoma. For almost all tumor suppressor genes, the entire gene region (exons and introns) were sequenced to facilitate CNV calling; for oncogenes, only exons were sequenced. For cases where the full gene is captured and sequenced, we separated the gene region into consecutive windows of 500 base pairs. This resulted in a panel of 1398 targets across 350 samples.

We applied CODEX2 to this data set and compare to CODEX. The number of Poisson latent factors in the background model is determined by the Bayesian information criterion (BIC) for both programs. The use of negative controls in estimating the background model allowed CODEX2 to be more robust to model tuning. For CODEX2, the number of latent factors had minimal effect on normalization and more generally on CNV detection, as only the normal samples were used to estimate the bias coefficient for each exon (Supplementary Table 7). In comparison, for CODEX, the number of CNV events decreased as the number of latent factors increased (Supplementary Table 7). Since the 108 genes are sparsely scattered across the genome, segmentation is performed within each gene separately. Furthermore, due to clonal heterogeneity and normal cell contamination, copy numbers may not be integers, and are assumed to be continuous and fractional to represent attenuated mean estimates of the genome mixtures. We categorize a CNA event to be high gain, gain, diploid, one-copy deletion, and two-copy deletion, if the profiled copy number is above 3.3, between 2.3 and 3.3, between 1.7 and 2.3, between 0.7 and 1.7, and below 0.7, respectively. Figure 4 shows the heatmaps of the segmentation results by CODEX and CODEX2. Each row corresponds to a sample, with the first 16 samples towards the bottom corresponding to the normal controls; each column corresponds to a target in the gene panel. In melanoma, somatic deletions of tumor suppressors (e.g., *PTEN*) and duplications of oncogenes (e.g., *BRAF*) are known to have high incidence rates [1]. From visual inspection of the heatmaps in Figure 4, we see that CODEX2 successfully retains these expected recurrent deletions and duplications, while CODEX, which does not make use of the negative control samples in fitting the background model, misinterprets the recurrent signals as a background latent factor.

**Figure 4.**
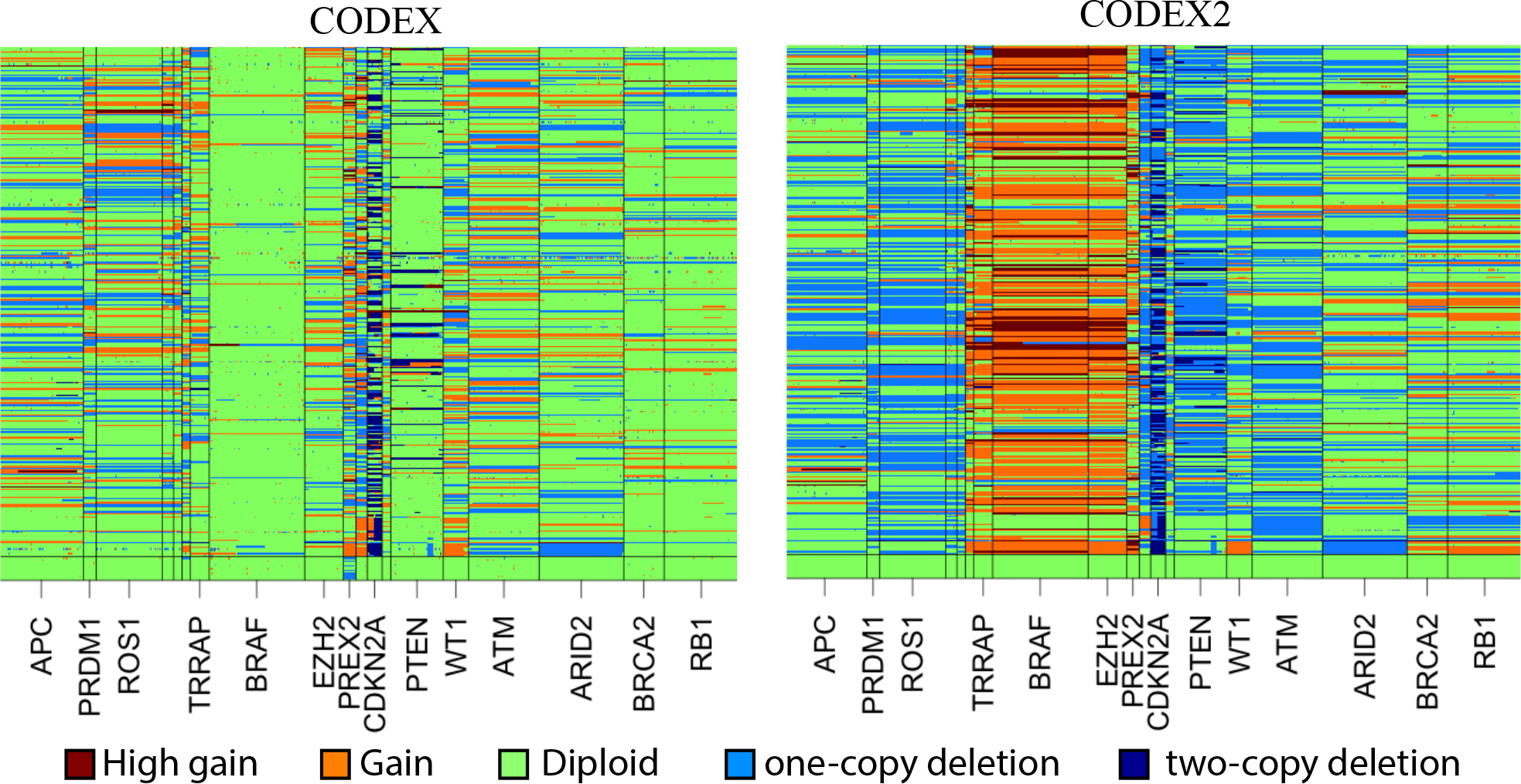
Heatmap of CODEX and CODEX2 normalization/segmentation results for the melanoma cohort. Each column is one target in the gene panel; each row is one sample, with the first 16 towards the bottom in the heatmap being normal. Profiled CNVs are categorized as high gain, gain, diploid (null), one-copy loss, and two-copy loss based on the estimated copy numbers. Only part of the oncogenes and tumor suppressors with greater than 30 targets are shown.

To rigorously evaluate CODEX2’s accuracy on this data, we compared the frequencies of the profiled gains and losses, that is, the proportion of samples in which a call is made, with frequencies from an independent melanoma patient cohort in TCGA [1]. Specifically, for each gene target, we plotted in Figure 5 the proportion of samples carrying a deletion (or duplication) in TCGA, versus this corresponding proportion in our current data set. CODEX2 achieves much higher concordance with TCGA results, with overall correlation across genes reaching 0.842 for deletions and 0.853 for gains, as compared to CODEX (correlation = 0.52 for deletions and 0.049 for gains). CODEX2 detects in these cell line samples a higher frequency of *BRAF* amplification and *CDKN2A* loss, as compared to the frozen-tissue derived TCGA results, which is not surprising due to the relative *in vitro* growth advantage of cells carrying these mutations. Based on the results by CODEX2, Garman et al. [27] further separated the cohort based on cancer subtypes and clinicopathological characteristics (responses to targeted and/or immunotherapy) and investigated the differences in mutational profiles between groups. The accurate profiling of CNVs in this data set enables unbiased downstream analysis.

**Figure 5.**
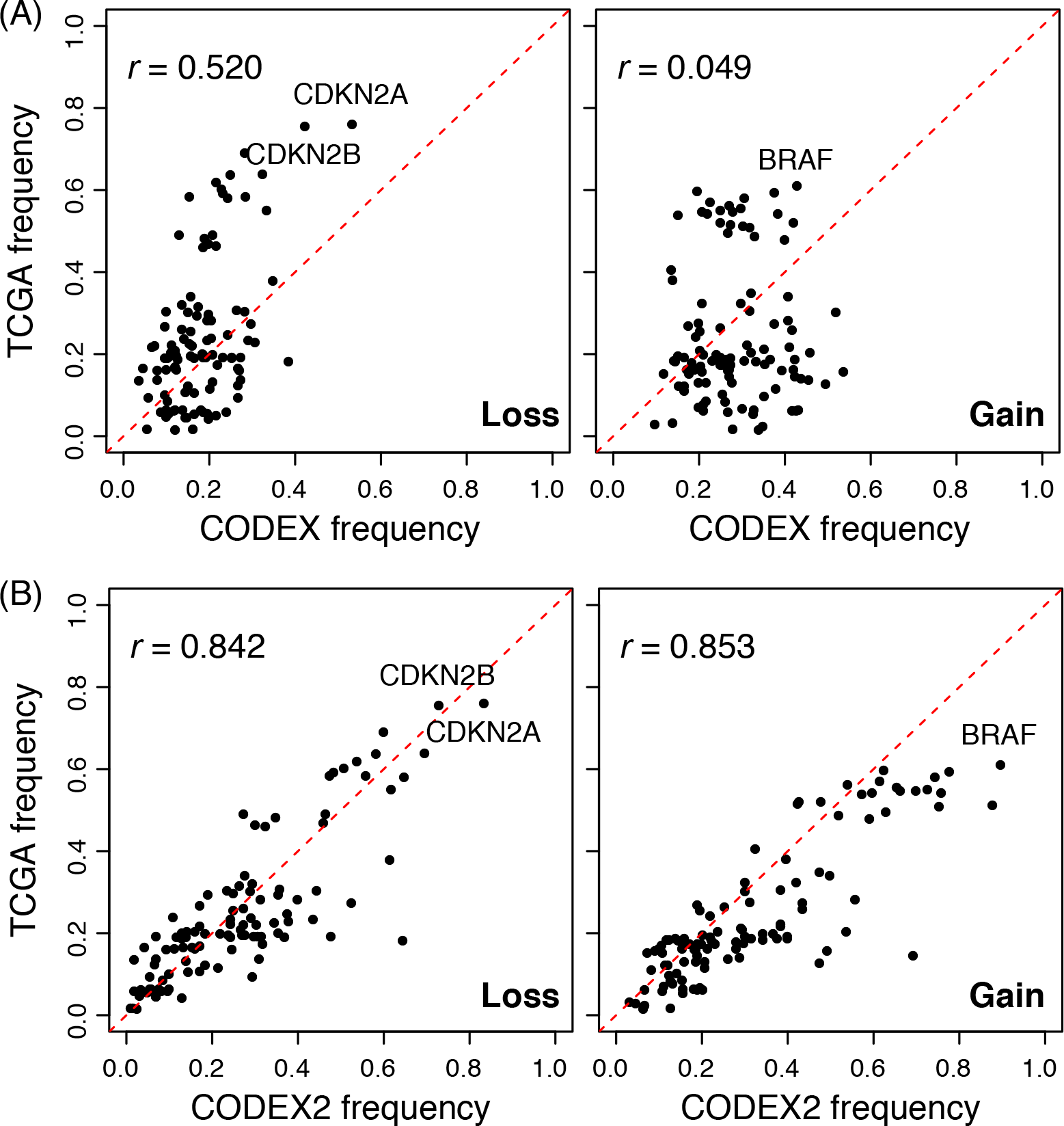
Assessment of profiled CNVs in the melanoma cohort with comparison to TCGA. CNVs are separated by states: losses on the left panels and gains on the right panel. Each dot corresponds to one gene in the targeted sequencing panel. CNV frequencies detected by (A) CODEX and (B) CODEX2 from the melanoma cohort is compared to the TCGA cohort with CODEX2 having drastically higher correlation.

### Performance assessment via spike-in studies

To understand how variables such as CNV length and population frequency affected the sensitivity of CODEX, CODEX2, and methods based on singular value decomposition (SVD, such as XHMM and CoNIFER), we conducted *in silico* spike-in studies. We started with the exon read depth data from chromosomes 16 to 22 in the 90 samples we analyzed from the 1000 Genomes Project, and applied filtering to remove putative existing CNVs. We then added, to the background count matrix under the null, heterozygous CNV signals of varying length, frequency, and degree of correlation with the first latent factor in the background model. See details of simulation setup in Methods.

As an illustration, Figure 6 shows a small subset of the CNV regions in the spike-in data with the ground truth, the post-normalization heatmap, and the CNV assignments across multiple methods, with the “null” regions containing no CNVs removed for easier visualization. The histograms in Figure 6 show the distribution of the normalized *z*-scores, with exons that harbor CNVs in red and exons within diploid regions in grey. We see that CODEX2 achieves clear separation of the deletions and diploids with the distributions centered on the expected values, log(1/2). The segmentations by CODEX and XHMM contain false negatives as well as false positives, especially in regions where common CNV signals reside, whereas improved normalization by CODEX2 allows almost perfect segmentation. Supplementary Figure 4 further shows the true latent background parameters (estimated using raw read depth without spike-in, which represents the null model that would be unobservable in a real data scenario) and the estimated parameters obtained by CODEX and CODEX2 on the “observed data” containing added CNVs. Our results show that while the sample loadings {***h***_1_,***h***_2_,***h***_3_} are consistent between CODEX, CODEX2, and the ground truth, the exon-specific factors {***g***_1_,***g***_2_,***g***_3_} estimated by CODEX are biased due to the inclusion of the mean CNV signals, reflecting the same trend in Supplementary Figure 5. CODEX2 corrects this bias through the use of the mixture model.

**Figure 6.**
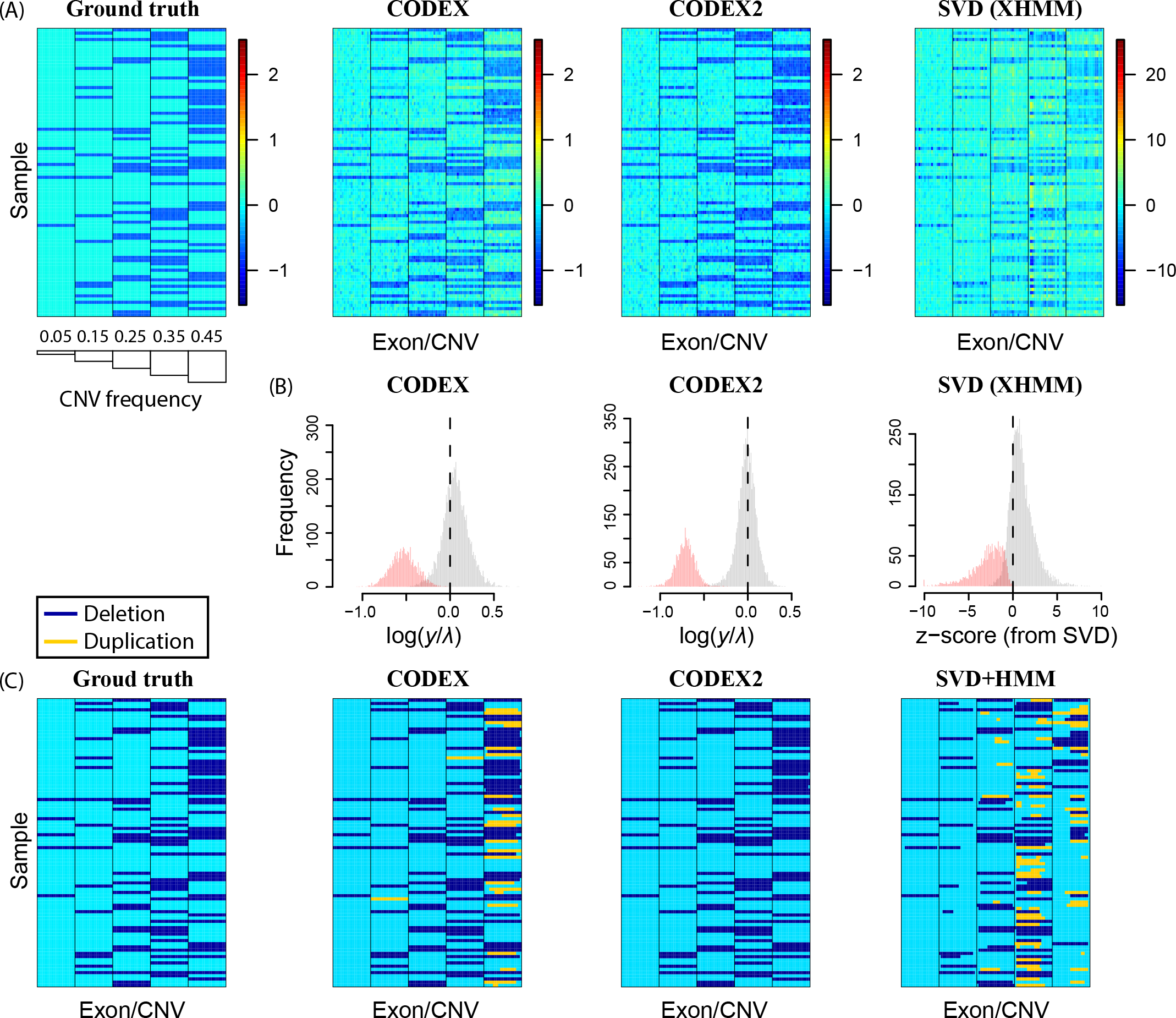
Performance assessment by spike-in experiment. CNV signals are *in silico* spiked in with increasing population frequencies. (A) Ground truth, and normalization results by CODEX, CODEX2, and SVD based method. (B) Histogram of normalized ***z**-*scores as 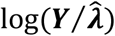 in CNV (red) and diploid (grey) regions. The ***z***-scores returned by XHMM are on a different scale with a much wider range and significant overlap between the null and non-null. (C) Segmentation results where CODEX2 has the highest concordance with the ground truth among the methods being compared.

We systematically compare the performance of CODEX2 against existing methods by spiking in deletions and duplications of length 5, 10, 20, and 40 exons with population frequency *p* ∈ {5%, 10%,…,95%}, repeating each simulation run 20 times. The precision and recall rates achieved by each method are shown in Figure 7, Supplementary Table 8, and Supplementary Table 9. Figure 7 shows how two variables, population frequency and degree of correlation with batch effect, impact the accuracy of methods. The results show that CODEX and SVD-based methods are sensitive to both variables, while CODEX2 maintains high accuracy across all frequencies and all degrees of correlation. CODEX2 has nearly perfect performance, whereas CODEX and SVD-based methods suffer from low sensitivity and specificity, especially for common CNVs with frequencies around 50%. We also investigated the effect of CNV length on performance and demonstrated that CODEX and SVD-based methods have lower sensitivity and specificity for longer CNVs, as compared to CODEX2 (Supplementary Figure 6, Supplementary Table 8).

**Figure 7.**
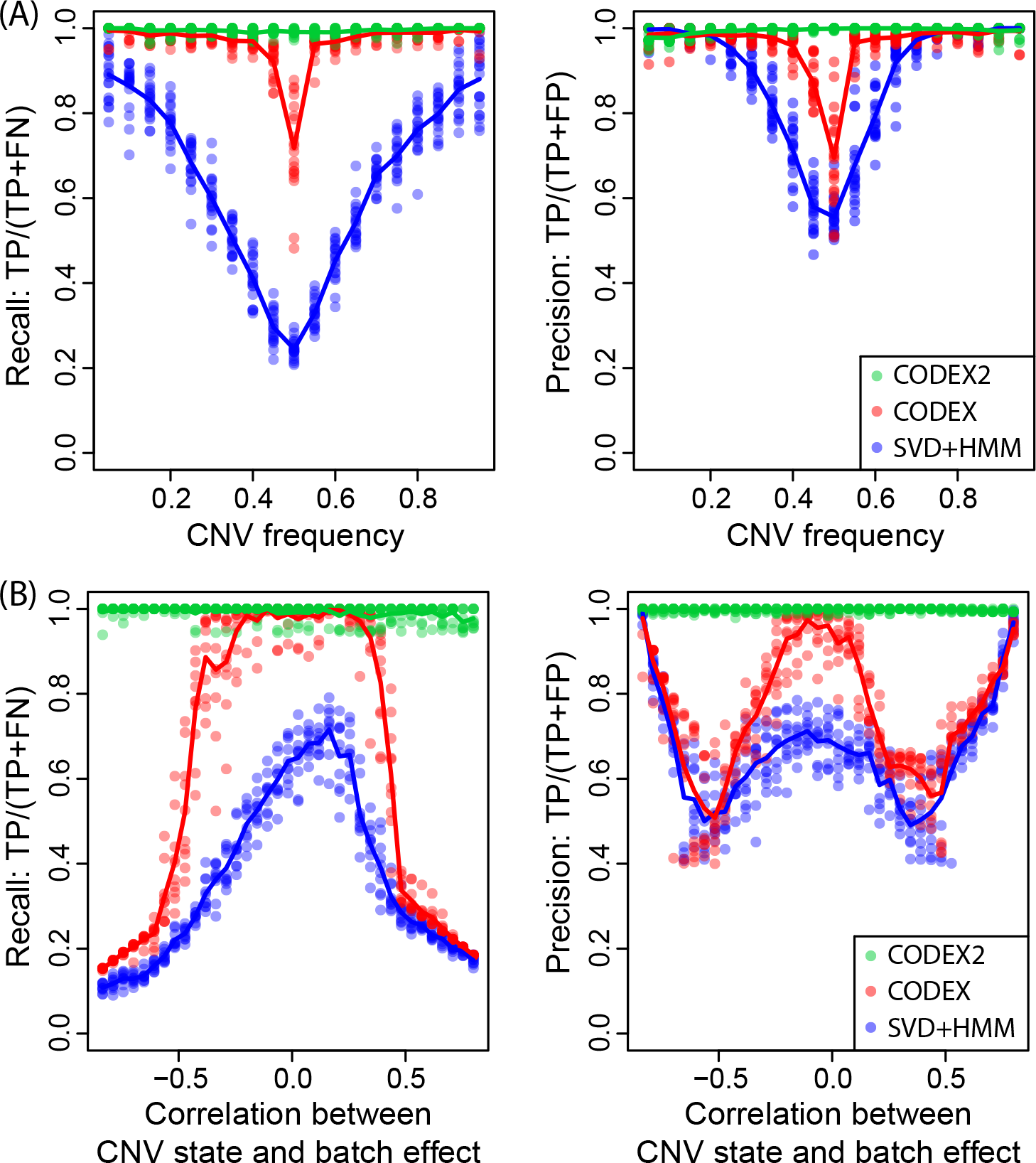
Assessment of precision and recall rates with different CNV frequencies and correlations between CNV state and batch effect. CODEX2 has nearly perfect performance compared to CODEX and XHMM. The latter two suffer from low recall and precision rates, especially for common CNV signals that are highly correlated with the batch effects.

We further studied the relationship between CNV carrier status and batch effects— 44 and 46 samples are sequenced by the Baylor College of Medicine and the Washington University Genome Sequencing Center respectively—and show that when they are highly correlated, CODEX suffers from low sensitivity, while CODEX2 is able to recover the CNV signals from the batch effect (Figure 7). Detailed results from an example run are shown in Supplementary Figure 7, where the spiked-in CNVs are highly correlated with the batch effect (i.e., most of the carriers are from one of the sequencing centers). CODEX estimates the latent factors assuming all exons are null, with the fitted regression line lying between the carriers and non-carriers, resulting in low sensitivity for true deletions in carriers and false positives as duplications in non-carriers. CODEX2, on the other hand, estimates the exon-specific latent factors in common CNV regions using the proposed EM algorithm and successfully separates the carriers from the non-carriers, which leads to clean down-stream segmentation. Here we show that normalization is of first-order effect.

We also performed additional simulations where we spiked in CNV signals to mimic those that are observed in genetically heterogeneous mixtures, such as cancer. Specifically, in each sample we added CNV signals as a mixture of *p*% cells with copy number *c* (sampled from a Gaussian distribution with mean 1 for heterozygous deletion) and (1 – *p*)% cells with copy number 2. If the copy number change event is clonal in the cancer sample, *p* is referred to as purity in cancer genomics study, which is the proportion of cancer cells out of the entire cell population; if the copy number change is subclonal, *p* is the cancer cell frequency for the CNV event. Our results show that CODEX2 is able to recover CNV signals with decent sensitivity and specificity for *p* as low as 30% (Supplementary Table 10). In the cancer genomics setting, we integrated CODEX2 with downstream software to create a stand-alone pipeline MARATHON [28] and there, demonstrated on a cancer phylogeny study of a neuroblastoma, breast cancer, and melanoma patient, with detection of somatic CNAs.

## Methods

### Background and overview

Multiple methods have been developed to recover CNV signals from experimental noise. VarScan2 [13], ExomeDepth [15], and ExomeCNV [16] control for baseline fluctuations in read coverage by relying on matched normal samples or building an optimized reference set. EXCAVATOR [14] adopts a median normalization approach for bias removal. It was soon realized that the magnitudes of the various sources of bias are sample-specific, and thus cannot be completely removed by normalizing to control samples or reference sets. This realization motivated the development of CLAMMS [19], where a reference panel is selected for each sample based on seven sequencing quality control metrics, as well as CoNIFER [10] and XHMM [11], which adopt SVD to estimate sample-specific backgrounds that can be more effective. CLAMMS, however, cannot be applied to WGS data or cancer samples and SVD is designed for capturing linear biases in continuous-valued observations. GC content has been shown to have a sample-specific, non-linear bias in sequencing data. Furthermore, read counts are not fit well by Gaussian models, even after transformation, due to their fluctuation over a very wide range. Our previous work, CODEX [12], adopts a Poisson latent factor model for count-based sequencing data and estimates a sample-specific background for each genomic position that incorporates nonlinear biases due to GC content, target-specific capture and amplification efficiency, and low-dimensional latent systemic artifacts.

We will start by giving an overview of the SVD-based methods by CoNIFER [10] and XHMM [11] and the Poisson latent factor model by CODEX [12]. We will discuss the limitations of existing models and the reason why they lack sensitivity for common CNVs. We will then describe the model for CODEX2, leaving algorithmic details to the Supplements.

### Review and reevaluation of existing methods

Denote ***Y*** as a *n × m* matrix of raw read depth, where *Y*_*ij*_ corresponds to the read depth for exon *i* ∈ {1, …, *n*} in sample *j* ∈ {1, …, *m*}.SVD-based methods CoNIFER [10] and XHMM [11] remove the strongest *K* SVD components,

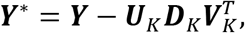

where ***U***_*K*_ and ***V***_K_ correspond to the top *K* left and right singular vectors, respectively, and ***D***_*K*_ corresponds to the diagonal matrix of the *K* largest singular values. Each column of ***Y**^*^* is column-standardized. The genome is then segmented by hidden Markov model (HMM) into ‘diploid’, ‘deletion’, or ‘duplication’ states.

This paper extends CODEX [12], which improves upon SVD-based approaches in several ways: CODEX adopts a Poisson model that more accurately models count data, and importantly, it explicitly models observable and measurable sources of bias, such as GC content and exon length, in addition to unmeasurable biases, due to unanticipated experimental variables, in the form of latent factors. In particular, GC content bias exhibits nonlinear patterns of variation across samples [7, 12], and thus CODEX uses a non-linear sample-specific function instead of a low-rank linear factor to capture this bias. Since an understanding of the CODEX model is integral to our ensuing discussion, we review it here. CODEX assumes that

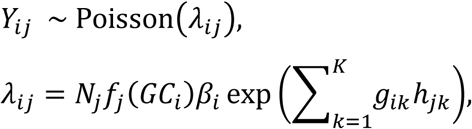

where *N*_*j*_ is the total number of reads in sample *j, f_j_*(⋅) is the GC content bias function for sample *j*, *GC*_*i*_ is the GC percentage of exon *i*, *β*_*i*_ is an exon-specific factor capturing multiplicative effects due to features such as exon length, and ***g**_k_* = {*g_1k_,.,g_nk_*}, ***h**_k_* = {*h_1k_, …, h_mk_*} are the *k*th (1 ≤*k ≤ K*) Poisson latent factors. The Poisson latent factors form a low-dimensional background model to capture unanticipated experimental variables, similar to the singular vectors in SVD-based normalization models [10, 11]. CODEX uses maximum likelihood to estimate the parameters of the null model, where ***g*** and ***h*** are assumed to be orthogonal to ensure identifiability (see Supplementary Algorithm 1 for details). For the remaining details on model selection, parameter estimation, and genome segmentation, see Jiang et al. [12].

These SVD- and factor-based methods, so far, all lack sensitivity for common CNVs. This is because common CNVs bias the estimation of the low-dimensional linear factors (***g***_*k*_ and ***h***_*k*_ in CODEX’s model and ***U**_k_*, ***V***_*k*_ in SVD-based algorithms), which, in CNV regions, capture part of the CNV signal. Therefore, subsequent removal of these factors also removes the signal for the CNVs. Supplementary Figure 5 shows a toy example illustrating this issue, focusing on one exon *i\** and assuming that *K =* 1. For this exon, CNV signals are *in silico* added at six population frequencies, creating six scenarios. For each scenario, simulated read counts ***Y***_*i*_ = {*Y*_*i1*_, …, *Y*_*im*_} are plotted against the sample factors ***h***_*1*_ = {*h_11_, …, h_m1_*}. The samples that carry the CNV are shown in red, the rest are shown in black. In each iteration of CODEX, Poisson log-linear regression is performed for ***Y***_*i*_ on ***h***_*1*_ to get *g_i1_,* the ‘loading’ for exon *i*. CODEX estimates *g*_*i1*_ assuming that the great majority of the samples are ‘nulls’, that is, they don’t carry CNVs, resulting in estimated background values shown in green. Ideally, the background curve should be fit to the control samples, the true ‘nulls’, as shown in blue. Yet in real data we don’t know which are the true carriers of this CNV, and the background curve is thus contaminated with signal, lying between the carriers and the non-carriers. The higher the incident rate of the CNV, the higher the contamination of the background by signal, and the closer the fitted green curve to the carriers rather than the non-carriers. This, as a result, leads to low sensitivity in detecting the true CNV carriers. The effect of population frequency on detection sensitivity by CODEX and SVD-based algorithms is also explored by simulation shown in Figure 6 and Figure 7.

### CODEX2 model and full-spectrum CNV detection

Figure 1 gives an overview of the two scenarios of CODEX2. To describe CODEX2, we first consider the simpler scenario where negative control samples are available. Without loss of generality, we consider a cohort comprised of both tumor and normal samples, the normal samples may not be matched to the tumor samples. Our goal is to identify CNVs present in the cancer samples but not in the normal samples. Duplications of oncogenes and deletions of tumor suppressors are commonly seen in cancer samples and have been reported to be associated with cancer. It is therefore crucial to detect somatic CNVs recurrent only in cancer samples with high sensitivity and specificity. Denote ***J***_*c*_ as the set of indices of the normal samples, which serve as negative control. Assuming that the normal samples are copy-number-neutral, the negative control samples are used by CODEX2 to estimate the exon-specific bias **β** and latent factors ***g*** to avoid attenuating and removing common CNV signals. Poisson regression is then to each cancer sample to obtain the sample loadings ***h***. The sample-specific background values are then computed using ***β***,***g,h*** and used in Poisson likelihood segmentation to identify CNVs. Refer to Supplementary Algorithm 2 for the detailed estimation procedure.

Now let’s consider the scenario where negative control samples are not available. We denote ***I**\** as a set of indices of the exons that harbor highly recurrent CNVs, the compliment of which are the indices of exons within the negative control regions. The set ***I**\** can be obtained based on prior knowledge (e.g., common deletions in tumor suppressors), from existing database (e.g., the Database of Genomic Variants), or empirically from a first-pass CODEX run. That is, if an exon lies within a common CNV region, CODEX will return a high standard deviation of the normalized *z*-scores across all samples for this exon - Supplementary Figure 5 shows that for a common CNV, the estimated null will be biased towards the alternative, especially when the incidence rate is high. Figure 1B shows an example on identifying germline CNVs from a population of samples (e.g., healthy individuals from the 1000 Genomes Project). For step 1 in Supplementary Algorithm 2, we no longer have a set of controls samples to directly estimate the exon-specific parameters. We propose an expectation-maximization (EM) algorithm embedded in our iterative parameter estimation procedure, where the missing data is the carrier status of the samples. Specifically, for each exon *i* ∈ ***I***\*,

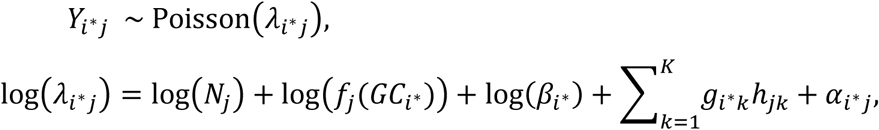

Where

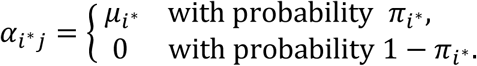

*π*_*i*\*_ is the incident rate for the CNV that span exon *i*,* and *μ*_*i*\*_ is the deviation from the null on the log scale, which can be either pre-fixed (e.g., log(1/2) for heterozygous deletion) or estimated by CODEX2. For simplicity, here we show the case where there is only one type of CNV event within the carriers. This can be easily extended to multiple subgroups with a model selection metric, which is enabled in the CODEX2 package. ***N, f(GC),*** and ***h*** can be using negative control regions shown in Supplementary Algorithm 3 step 1 and 2. We adopt an EM algorithm to estimate the unknown parameters 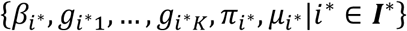with missing data:

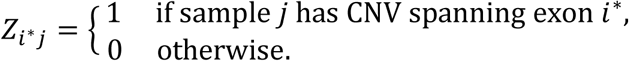

Refer to Supplementary Algorithm 3 for implementation details.

CODEX2 offers the choice of Akaike information criterion (AIC), Bayesian information criterion (BIC), and variance reduction to determine the optimal number of latent factors. In CODEX and SVD-based methods (XHMM and CoNIFER), the number of latent factors is a critical model tuning parameter that affects normalization and segmentation results. It is often not clear whether to use the AIC, BIC, or simple visual examination of the scree plot, and this arbitrariness plagues all methods that rely on SVD, PCA, or factor models. By using negative controls to guide the estimation of the background model, CODEX2 is less sensitive to the number of latent factors (Supplementary Table 7), thus giving results that are easier to reproduce.

For segmentation, CODEX2 adopts the same Poisson likelihood ratio based approach as CODEX. Please refer to Jiang et al. [12] for details. For targeted sequencing, where a smaller pre-selected panel of targets are sequenced, the normalization model is exactly the same; the segmentation is performed within each gene separately. For WGS, user-defined consecutive bins can be treated as ‘targets’ in the WES setting, with normalization and segmentation procedures carried out in the same fashion. Refer to Zhou et al. [6] for details.

### Simulation setup

We start with the exonic read depth data of chromosomes 16 to 22 from the 90 samples we analyzed from the 1000 Genomes Project, and apply filtering to remove putative existing CNVs. Specifically, the filtering step removes all exons that: (i) are called to harbor CNV events by XHMM, EXCAVATOR, CODEX, or CODEX2, (ii) overlap with duplication and deletion reported in the Database of Genomic Variants, (iii) don’t pass quality control procedure by CODEX (median coverage between 40 and 4000, length between 30 and 2000 base pairs, mappability less than 0.95, GC content between 30% and 70%), (iv) have standard deviation of normalized z-scores across samples above 0.3, or maximum of normalized z-scores above 0.8, or minimum normalized z-scores below - 0.8 across all samples. This way we are left with 4035 ‘null’ exons that are CNV-free across 90 samples. Treating this filtered count matrix as background, we fit the background model of CODEX, with the estimated parameters as ground truth. The optimal number of latent factors is 3 by AIC, BIC and variance reduction (Supplementary Figure 4) and is kept the same for subsequent analysis for CODEX2, CODEX, and SVD-based method. We then add, to this background count matrix, heterozygous CNV signals of varying length, frequency, and degree of correlation with the first latent factor in the background model. In more detail, we spike in heterozygous deletions, of varying lengths and population frequencies, by reducing the raw depth of coverage for exons spanned by the CNV from *y* to *y* × *c*/2, where *c* is sampled from a normal distribution with mean 1 and standard deviation 0.1. Note that heterozygous deletions with frequency *p* in the population have exactly the same detection accuracy as duplications with frequency 1 – *p*, since all copy number events are defined in reference to a population average. To confirm, we also spike in copy number gains with varying population frequencies where *c* is sampled from a Gaussian distribution with mean 3. To assess performance in cancer genomics setting where the samples are heterogeneous, we further carry out spike-in studies where CNV signals are added as a mixture, with *p*% tumor cells having copy number *c* and (1 – *p*)% normal cells having copy number 2.

### Discussion and conclusions

A limitation shared by all existing CNV detection methods, highlighted by multiple independent benchmarking studies, is the lack of sensitivity for common variants. Similarly, in our experience applying CODEX [12] to WES and targeted sequencing of tumor samples, we easily detect sporadic aberrations but miss highly recurrent aberrations. To meet the widespread demand for improved CNV detection, we develop in this paper a new method, CODEX2, to remove technical noise and improve CNV signal-to-noise ratio for all sequencing platforms including WES and targeted sequencing. CODEX2 builds on our existing method CODEX [12] with a significant improvement of sensitivity for common variants, thus allowing full-spectrum CNV detection. CODEX2 can be applied in either the setting of a case-control analysis in which the goal is to detect CNVs that are enriched in the case samples, or when the goal is simply to profile all CNVs.

We have benchmarked CODEX2 extensively against existing methods. In the first evaluation, we reanalyzed WES data of the HapMap samples from the 1000 Genomes Project, for which a set of experimentally validated CNV calls from microarrays and other sources could be used to assess performance. Our results demonstrate that CODEX2 has markedly improved sensitivity and specificity over existing methods. The improvement for calling of common variants is the most substantial, from 40% recall rate to >80% recall rate, in two out of the three validation sets used, while simultaneously improving precision from 60% to 90%. In the second evaluation, we applied CODEX2 to targeted sequencing data of melanoma cell lines, PDX, and tumor samples, in which CODEX2 detects CNVs with recurrence rates that are highly concordant with those obtained from TCGA. Finally, we performed extensive simulations benchmarking existing methods and elucidated how key variables, such as population frequency, influence detection sensitivity. Together, these results establish the improved accuracy of CODEX2 over existing state-of-the-art approaches, and the utility of the software under varying study designs.

Under a different context, for the detection of differential expression in RNA sequencing data, Risso et al. [29] proposed the normalization method "removing unwanted variation” (RUV) which is based on a factor model that relies on sets of control genes or samples for estimation. CODEX2 resembles RUV but has distinguishing features. First, RUV is designed to be used in a case-control setting. CODEX2 can be applied with and without negative control samples. Second, when estimating the latent factors that correspond to the control genes or samples, RUV adopts SVD with Gaussian assumption, whereas CODEX2 uses Poisson generalized linear modeling to achieve better fit for low counts. Furthermore, CODEX2 directly models the GC content bias, which cannot be fully captured by a linear principal component [7, 12], as well as library size and exon length, all of which can be directly quantified.

CODEX2 normalizes the read depth data for CNV detection via a Poisson latent factor model, which can be well adapted to other settings within the genomics domain. Lee et al. [30] applies a similar Poisson factor approach to the non-normalized microRNA sequencing data. Chen et al. [31] estimates allele-specific copy number under tumor-normal setting using the Poisson latent factor model to remove biases and artifacts that cannot be fully captured by comparing to the normal.

CODEX2 has been integrated into a pipeline, iCNV, to detect CNV using both sequencing (WGS, WES, and targeted) and microarray data [6]. For cancer genomics studies, CODEX2 has been integrated into a pipeline, MARATHON, to infer both germline and somatic copy number changes and reconstruct tumor phylogeny [28]. Sequencing capacity has increased exponentially over the past few years. This tremendous amount of data opens up great opportunities and yet at the same time precautions should be made with regard to running efficiency and method scalability. CODEX2 processes each chromosome within each batch separately and can thus be highly parallelized. With increasing sequencing capacity, and increasing need to profile CNVs as a non-negligible source of genetic variation, we believe that CODEX2 can be a useful tool for the genetics and genomics community.

## Abbreviations

CNV: Copy number variation
WGS: whole-genome sequencing
WES: whole-exome sequencing
SVD: singular value decomposition
TCGA: the Cancer Genome Atlas
AIC: Akaike information criterion
BIC: Bayesian information criterion
EM: expectation-maximization
RUV: remove unwanted variation

## Declarations

### Ethics Approval and consent to participate

Not applicable.

### Consent for publication

Not applicable.

### Availability of data and material

CODEX2 is an open-source R package available at https://github.com/yuchaojiang/CODEX2 with license GPL-2.0.

## Authors’ contributions

NRZ initiated and envisioned the study. YJ and NRZ formulated the model. YJ developed and implemented the algorithm. YJ, RW, EU, and NRZ performed simulation studies and the HapMap analysis. YJ, INA, KLN, and NRZ conducted the melanoma analysis. YJ and NRZ wrote the manuscript, which was edited by KLN. All authors read and approved the final manuscript.

## Acknowledgements

We thank Bradley Garman and Bradley Wubbenhorst for providing the case-control melanoma dataset from samples collected and studied at the Wistar Institute in the laboratory of Dr. Meenhard Herlyn, led by Dr. Clemens Krepler. We also thank Xiaoqi Geng, Xiangwei Zhong, and Dr. Jingshu Wang for helpful comments and discussions.

